# Ethylene mitigates waterlogging stress by regulating glutathione biosynthesis-related transcripts in soybeans

**DOI:** 10.1101/252312

**Authors:** Yoonha Kim, Chang-Woo Seo, Abdul Latif Khan, Bong-Gyu Mun, Raheem Shahzad, Jung-Woo Ko, Byung-Wook Yun, In-Jung Lee

## Abstract

Waterlogging stress is a restrictive factors in soybean productivity worldwide. Plants utilize various physio-chemical changes to mitigate waterlogging stress. In current study, the regulatory roles of seven kinds of plant growth regulators, including abscisic acid (ABA), ethylene (ethephon, ETP), gibberellins (GA4), indole-3-acetic acid (IAA), kinetine (KT), jasmonic acid (JA) and salicylic acid (SA), were determined for soybeans under waterlogging conditions. Based on the results, the donor source of ethylene was selected and its role was further examined regarding physiochemical alteration and glutathione biosynthesis-related transcripts through application of exogenous ETP. ETP application mitigated waterlogging stress and significantly improved the efficiency of photosynthesis and increased bioactive GA4 contents compared to that of untreated plants. Element and amino acid contents among the treatments were significantly different. Total elements and amino acid contents were increased in 100 μM ETP-treated soybean plants. ETP application induced adventitious root initiation, improved root surface area, and significantly increased glutathione transferases expression and glutathione relative to that of non-ETP treated soybean plants. Finally, 100 μM-ETP application induced up-regulated protein content and glutathione s-transferase DHAR2 as compared to that of soybeans under waterlogging-conditions only. ETP could induce various biochemical and transcriptional modulations that strengthen plant growth and mitigate waterlogging stress.

**Highlight:** Ethylene application to soybean plants after waterlogging up-regulates glutathione transferase genes. Higher glutathione activity, as well as increased glutathione s-transferase DHAR2 protein content was induced to scavenge reactive oxygen species.

## Introduction

Flooding stress negatively influence most of the plants physiological functions which leads to poor photosynthesis, hormonal imbalance and reduced nutrients uptake and resulted in stunted growth and yield (Valliyodan *et al*., 2017). According to NASA, approximately 17 million km^2^ of land area worldwide have been exposed to flooding conditions. (Perata *et al*., 2011). However, among flooding conditions, waterlogging (WL) is a more common problem than submergence. WL stress in field conditions occurs for several reasons, including overflow of rivers and heavy rainfall (Kim *et al*., 2015).

In South Korea, soybeans are not only regarded as an important field crop because of their higher nutritional value, but also because it is a higher income crop than paddy-field crops, such as rice (Koo *et al*., 2014). According to Park *et al.* (2015), the Rural Development Administration (RDA) in South Korea stated that soybean field and rice paddy field income was $645/10 a and $544/10 a, respectively in 2012. Thus, profit per field unit was 15.6% higher for soybean fields in comparison with that of rice paddy fields. For this reason, to increase farm income the RDA in South Korea strongly recommends soybean cultivation in paddy field (only 16.9% of land area could be used for soybean crop cultivation) (Park *et al*., 2015). Generally, soybeans are a flood-intolerant crop, for which growth and yield are negatively affected by WL stress. According to Nguyen *et al.* (2012), soybean yield was 17% (vegetative stage) and 57% (reproductive stage) less when soybean plants were exposed to WL stress conditions as compared to that of non-stress conditions. Thus, several soybean breeders have been developing tolerant varieties against WL stress or have been focused on identification of tolerance mechanisms under WL conditions (Reyna *et al*., 2003; Nishiuchi *et al*., 2012). In particular, our research team recently reported the physiological differences between a WL-tolerant soybean variety and a WL-susceptible soybean variety, and confirmed significant alteration in different endogenous hormones levels (abscisic acid: ABA, ethylene: ET, gibberellins: GA, salicylic acid: SA, jasmonic acid: JA) in the contrasting soybean lines, as well as differences in antioxidant activities and root architecture, such as lateral roots and aerenchyma cells (Kim *et al*., 2015). Among several physiological responses, our research team focused on different the ET level because ET is biosynthesized by the short-haul pathway in comparison with other plant hormones and is produced by 1-aminocyclopropane-1-carboxylic acid (ACC) oxidation. Therefore, oxygen is a main component of the production of ET (Arc *et al*., 2013). Consequently, plants cannot produce enough ET without adequate oxygen supply. However, ET production is significantly increased under WL conditions because of the enrichment of ACC synthase (Dat *et al*., 2004). Furthermore, ET is known to mitigate WL stress via development of aerenchyma cells with crosstalk among ABA, GA, indole-acetic-acid (IAA) and kinetin (KT) (Fukao and Bailey-Serres, 2008; Shimamura *et al*., 2014). In addition to crosstalk among ABA, GA, IAA, and KT, ET can interact with JA and SA to induce development of adventitious roots and aerenchyma cells (Brodersen *et al*., 2005; Yang *et al*., 2013).

Therefore, we decided to conduct a follow-up experiment to investigate whether exogenously applied plant growth regulators (PGRs) could induce WL tolerance. Thus, we treated to soybean plants under WL conditions to different PGRs; then, we looked for beneficial effects among PGRs. Thus, we conducted further experiments to identify physiological responses against WL stress via various experimental technologies.

## Materials and methods

### Selection of proper plant hormone (EPI)

To obtain more concrete results, the current study composed two experiments. Experiment I (EP I) was conducted to screen for the appropriate plant growth regulators in soybean plants to enhance resistance to WL stress conditions, and experiment II (EP II) was conducted to identify the physiological and biochemical mechanisms involved with plant hormone application during WL-stress mitigation. Thus, we applied different kinds of PGRs including ABA, ethylene (ethephon, ETP), GA_4_, IAA, KT, JA, and SA to soybean plants grown under WL-stress conditions.

## EP I

### Plant growth condition and PGR application

We used the Daewon variety (*Glycine max* L.) as the plant material because it is the most common variety and is broadly grown in South Korea. The seeds were donated from the National Institute of Crop Science, RDA, South Korea. The seed surfaces were sterilized with 70% ethanol and then thoroughly rinsed with autoclaved double distilled water. Seeds were sown into plastic trays (50 holes, 40 cm × 20 cm) filled with autoclaved horticultural soil (Tobirang, Baekkwang Fertility, South Korea) and they were then grown in green houses located in Kyungpook National University, Daegu, South Korea. Uniformly grown seedlings were transferred into plastic pots (6 holes, 455 mm × 340 mm × 180 mm) at 10 days after germination. Soybean plants (V2 stage) were subjected to WL stress conditions for two weeks (14 days). PGRs were applied to soybean seedlings 1 h after WL treatment. Detailed information about PGRs is given in Supplementary Table. 1.

**Table 1.** Protein information for soybean plants exposed to WL stress for 10 days.

| Spot No. | MW | PI | Protein name | Score | Accession No. |
| --- | --- | --- | --- | --- | --- |
| 615 | 53.03 | 6.04 | ribulose-1,5-bisphosphate carboxylase/oxygenase large subunit, partial (chloroplast) | 225 | YP_538747 |
| 616 | 51.57 | 6.04 | ribulose-1,5-bisphosphate carboxylase/oxygenase large subunit, partial (chloroplast) | 212 | SBO07506 |
| 1002 | 17.01 | 5.03 | trypsin inhibitor A | 119 | XP_003532237 |
| 1104 | 20.77 | 5.03 | glutathione s-transferase DHAR2 | 116 | AJE59632 |
| 1611 | 54.88 | 4.94 | ribulose-1,5-bisphosphate carboxylase/oxygenase large subunit, partial (chloroplast) | 204 | CAB08877 |
| 170 | 66.80 | 4.94 | ribulose-1,5-bisphosphate carboxylase/oxygenase large subunit | 216 | CAB08877 |
| 4101 | 18.55 | 5.91 | glycoprotein | 148 | NP_001241536 |

Accession number (GI number), MW: molecular weight; PI: isoelectric point

### Evaluation of resistance to WL stress

To evaluate mitigation effects on WL stress, we collected growth attributes, including plant height, chlorophyll content, and chlorophyll fluorescence during WL conditions and after WL conditions. Chlorophyll content was measured using a chlorophyll content meter (CCM-300, Opti-Sciences, USA) and chlorophyll fluorescence data were recorded using a chlorophyll fluorometer (OS5p+, Opti-Sciences, USA). The selection of proper plant hormones to enhance the resistance of soybeans to WL stress was conducted three times under greenhouse conditions and each experimental set was composed of three replications.

## Experiment II

### Plant growth conditions and PGRs application

We confirmed stress mitigation effects of EP I. According to our experiment, application of ETP enhanced resistance to WL stress in soybeans relative to other PGR treatments. Thus, we applied three different concentrations of ETP to soybean plants to identify physiological and biochemical responses against WL stress.

### Plant growth conditions and ETP application

We used the same seeds, soil, and pots for EP I seed germination, but seeds were grown in a growth chamber [Day 30□ (14 h) / Night 22□ (10 h), relative humidity 70%, light intensity 1000 μmol m^-2^ s^-1^] to collect accurate data. After seed germination, the uniformly grown soybean seedlings were selected and transplanted into 6 holed pots (same size as with EPI) and maintained in the growth chamber. WL stress was applied to soybean plant at the V2 stage and water level was maintained at 10 cm–15 cm above the soil surface for 10 days. Three different concentrations of ETP were applied to soybean shoots 1 h after exposure to WL stress (Supplementary Table 1).

### Analysis of chlorophyll content and chlorophyll fluorescence

Chlorophyll content and fluorescence data were collected at 5, 10, and 15 days after WL stress. The chlorophyll content and fluorescence data were measured using the same methods that were described for EP I. The data were collected three times and each replication was composed of seven plants (n = 7).

### Endogenous hormones analysis

To analyze endogenous GA contents, we harvested the whole plant at 5, 10 and 15 days after WL conditions. Plant samples were immediately frozen in liquid nitrogen followed by freeze drying (ISE Bondiro freeze dryer, Operon, South Korea) for 5 days. Thoroughly dried plant samples were ground into very fine powder form after which they were used for GA analysis. We used 0.3 g dried samples used for GA analysis and followed the same analysis protocol described in Kim *et al.* (2015). Endogenous GA content was analyzed by GC-MS-SIM. In particular, endogenous GA_4_, GA_9_, and GA_34_ content were calculated from the peak area ratios of 284/286, 298/300, and 506/508, respectively. Data were collected in three times replications (n = 3) and the DMRT test was conducted to compare among treatments. Conditions for each instrument for hormone analysis are given in Supplementary Table 2.

### Macro mineral contents

The nutrient contents were analyzed according to the methods describes by (Jin and Zhu (2000). Briefly, 0.1 g of sample was weighted in the vials, extracted with HNO_3_ + H_2_O_2_ solution, and degraded by microwave. The solution was further diluted by adding double distilled water and then it was used to analyze the compounds. Carbon (C), hydrogen (H), oxygen (O), nitrogen (N), and sulfur (S) contents were analyzed using an elemental analyzer (Flash 2000, ThermoFisher, USA). Phosphorus (P), potassium (K), calcium (Ca), magnesium (Mg), and iron (Fe) were measured by ICP spectrophotometer after samples were moved into an elemental analyzer machine (Optima 7300DV, PerkinElmer, USA).

### Protein sample preparation

Soybean plant samples were washed twice with ice-cold PBS (in molecular cloning), sonicated for 10 s by Sonoplus (Bandelin electronic, Germany), and homogenized directly by mortor-driven homogenizer (PowerGen125, Fisher Scientific, USA) in sample lysis solution composed of 7 M urea, 2 M thiourea containing 4% (w/v) 3-[(3-cholamidopropy) dimethyammonio]-1-propanesulfonate (CHAPS), 1% (w/v) dithiothreitol (DTT), 2% (v/v) pharmalyte, and 1 mM benzamidine. Occasionally bead beater was used for lysis of rigid cells. Proteins were extracted for 1 h at room temperature with vortexing. After centrifugation at 15,000 × *g* for one h at 15°C, insoluble material was discarded, and the soluble fraction was used for two-dimensional gel electrophoresis. Protein concentration was assayed by the Bradford method (Bradford, 1976).

### 2D PAGE

IPG dry strips (4-10 NL IPG, 24 cm, Genomine, Korea) were equilibrated for 12 h–16 h with 7 M urea, 2 M thiourea containing 2% 3-[(3-cholamidopropy)dimethyammonio]-1-propanesulfonate (CHAPS), 1% dithiothreitol (DTT), and 1% pharmalyte and loaded with 200 μg of the sample. Isoelectric focusing (IEF) was performed at 20°C using a Multiphor II electrophoresis unit and EPS 3500 XL power supply (Amersham Biosciences, UK) following the manufacturer’s instruction. For IEF, the voltage was linearly increased from 150 to 3,500 V during 3 h for sample entry followed by constant 3,500 V, with focusing complete after 96 kVh. Prior to the second dimension, strips were incubated for 10 min in equilibration buffer (50 mM Tris-Cl, pH 6.8 containing 6 M urea, 2% SDS, and 30% glycerol), first with 1% DTT and second with 2.5% iodoacetamide. Equilibrated strips were inserted onto SDSPAGE gels (20 × 24 cm, 10 %–16%). SDS-PAGE was performed using the Hoefer DALT 2D system (Amersham Biosciences, UK) following the manufacturer’s instruction. 2D gels were run at 20°C for 1,700 Vh. Then, 2D gels were Colloidal CBB stained as previously described (Oakley *et al*., 1980), but the fixing and sensitization step with glutaraldehyde was omitted.

### Image analysis

Quantitative analysis of digitized images was conducted using the PDQuest (version 7.0, Bio-Rad, USA) software according to the protocols provided by the manufacturer. Quantity of each spot was normalized by total valid spot intensity. Protein spots were selected for significant expression variation that deviated over two-fold in its expression level compared with that of the control or normal sample.

### Identifications of Proteins

For protein identification by peptide mass fingerprinting, protein spots were excised, digested with trypsin (Promega, Madison, WI), mixed with ‐cyano-4-hydroxycinnamic acid in 50% acetonitrile / 0.1% TFA, and subjected to MALDI-TOF analysis (Microflex LRF 20, Bruker Daltonics, USA) (Fernandez *et al*., 1998). Spectra were collected from 300 shots per spectrum over m/z range 600 – 3000 and calibrated by two-point internal calibration using Trypsin auto-digestion peaks (m/z 842.5099, 2211.1046). Peak list was generated using Flex Analysis 3.0. Threshold used for peak-picking was as follows: 500 for minimum resolution of monoisotopic mass, 5 for S/N. The search program MASCOT, developed by the Matrixscience (http://www.matrixscience.com) was used for protein identification by peptide mass fingerprinting. The following parameters were used for the database search: trypsin as the cleaving enzyme, a maximum of one missed cleavage, iodoacetamide (Cys) as a complete modification, oxidation (Met) as a partial modification, monoisotopic masses, and a mass tolerance of ± 0.1 Da. PMF acceptance criteria was used for probability scoring.

### Root phenotype

We used the same soybean variety for root phenotyping. The sterilized soybean seeds were propagated in 6-hole pots (455 mm × 340 mm × 180 mm), which contained thoroughly washed decomposed granite soils (overall size was 7 mm–9 mm) to reduce root sample loss. To prevent soil drying, sufficient water was supplied in the morning (8 AM – 9 AM) and evening (18 PM – 19 PM). When soybean plants reached the V2 stage, we produced the WL condition for soybean plants for 15 days. During the WL period, water level was maintained daily at 10 cm–15 cm above the soil surface and the root samples were collected at 5-day interval after WL treatment began until day 15. Decomposed granite soil was carefully removed from pots and then root samples were washed with distilled water two times. Clean root samples were photographed using a digital camera (COOLPIX A, Nikon, Japan) in a mini studio (W 70 cm × L 100 cm). The image data was analyzed by the Flower Shape Analysis System (www.kazusa.or.jp/phenotyping/picasos, Tanabata *et al*., 2010) to determine root surface area (RSA).

### Antioxidant activity and mRNA expression level (EPII)

To measure stress response of soybean plants, we used the same plant material for an experiment of RSA. At the V2 stage, the WL condition was applied to soybean plants for 2 days. We collected plant samples at 1-day intervals and the same experiment was conducted three times to increase reliability. The fresh leaf and root samples were used to determine glutathione (GSH) and glutathione reductase (GR) activity. Briefly, 100 mg of fresh leaf samples was homogenized in buffer containing 50 mM Tris HCl (pH 7.0), 3 mM MgCl_2_, 1 mM EDTA, and 1.0% PVP. The homogenized samples were then centrifuged at 10,000 rpm for 15 min at 4°C. We used the Bradford assay for quantification of total protein content (Bradford, 1976). The reduced GSH was estimated following the protocol of Ellman (1959). The homogenate was collected by grinding leaf samples following the addition of 3 ml of 5% (v/v) trichloroacetic acid. The supernatant (0.1 ml) was decanted into a tube containing 3 ml of 150 mM NaH_2_PO_4_ (pH 7.4). Subsequently, 500 μl of 5.5´-dithiobis (2-nitrobenzoic acid) (DTNB; 75.3 mg of DTNB dissolved in 30 ml of 100 mM of phosphate buffer, pH 6.8) was added to the suspension and then it was incubated at 30 ± 2 °C for 5 min. The absorbance was measured at 412 nm. Thus, GSH contents were estimated by comparison with the standard curve. The GR activity was measured by the previously described protocol of Garlberg and Mannervik (1985). A total 150 μl of enzyme was reacted with 1 ml of reaction mixture containing 1 mM EDTA, 3 mM MgCl_2_, 0.5 mM oxidized glutathione, 0.1 M HEPES pH 7.8, and 0.2 mM NADPH. Activity of GR was measured by NADPH oxidation and was monitored by the decreased absorbance at 340 nm for 2 min.

### Statistical analysis

The experiments (EPI, EPII) were conducted three times with three replications in growth chamber and greenhouse environments. Experiments of antioxidant activity and its mRNA expression level, SNO‐ and SNO-related gene expression levels were conducted two times with three replications under greenhouse conditions. An analysis of variance (ANOVA) was used to evaluate significant differences (*P < 0.05*) in treatments, periods, replications, and treatments by periods. Comparisons between treatments was conducted by Duncan’s Multiple Range Test (DMRT) and SAS 9.1 software was used for statistical analysis.

## Results

### Experiment I

#### Plant growth characteristics with and without waterlogging (WL) stress

To evaluate PGR effects, we monitored plant height during and after WL stress. Soybean plant height showed non-significant differences among PGR treatments. As we anticipated, plant height was significantly increased in the GA_4_ treatment at 7, 14, 21, and 28 days after treatment (DAT; Fig. 1). In spite of increased plant height in the GA_4_ treatment, plants did not show WL resistance to lodging (data not shown). Despite the GA_4_ treatment, wherein increased plant height was observed in the ETP and KT application as compared to the WL-only treatment, treatments in which IAA, SA, and MeJA was applied to soybean plants did not show differences in comparison with WL-only treatment (Fig. 1). In particular, soybean plants died after 14 days of WL in the 100 μM ABA treatment (Fig. 1). To determine effects on the ability of the photosynthetic apparatus, we measured chlorophyll content and chlorophyll fluorescence after ETP and KT application. The chlorophyll content and chlorophyll fluorescence were not different when compared with that before WL treatment (Fig. 2). Chlorophyll content was significantly decreased in all WL-treated soybean plants. However, ETP‐ and CK-treated soybean plants had increased chlorophyll contents as compared to WL plants during the experiment periods (Fig. 2A). Moreover, in comparison between the ETP and KT application, chlorophyll concentration was higher with the ETP application (Fig. 2A). Very similar tendencies were observed for the determination of chlorophyll fluorescence. Chlorophyll fluorescence of hormone-treated soybean plants showed increased levels (45% – 50% increased at 28 DAT) as compared to that of WL-only plants. In particular, the chlorophyll fluorescence value in ETP-applied plants was higher than that of plants with KT application (Fig. 2B).

**Fig. 1.**
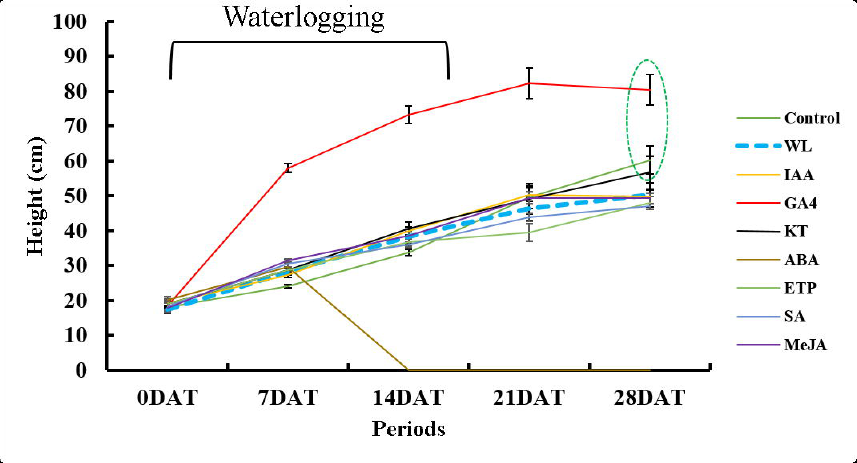
Soybean plant height during and after WL treatment. Waterlog condition was maintained for 14 days. In the figure, WL indicated only WL treatment thus, it was marked as dotted linear and green dotted circle revealed significant difference as compared to WL treatment. Data was collected by 3 times replication and it showed the average ± standard error (*n = 10*).

**Fig. 2.**
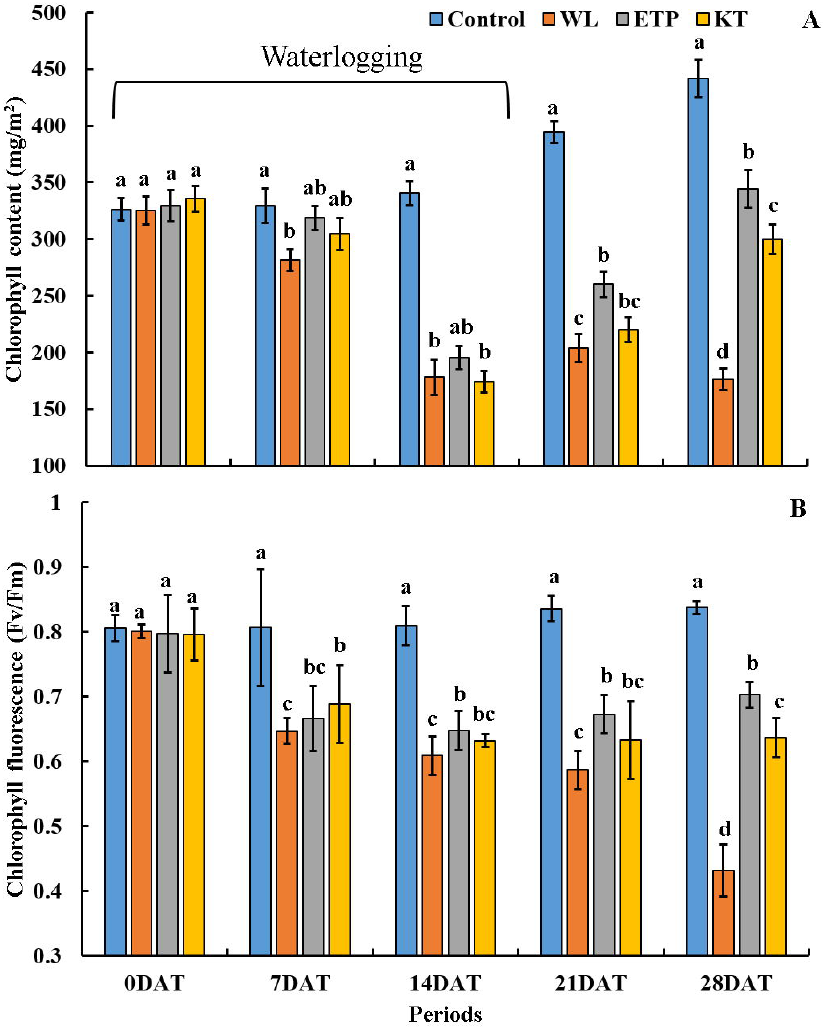
Changes in chlorophyll content during and after WL treatment. Waterlogging conditions were maintained for 14 days. In the figure, WL indicates WL-only treatment. Different letters in figure indicate significant differences at *P < 0.01* and data were analyzed by Duncan’s multiple range test (DMRT). Data were collected for three replications and the average ± standard error are shown (*n = 10*).

### Experiment II

#### Effects of ETP application on plant growth characteristics with and without WL stress

To obtain more detailed information related with stress mitigation, we applied three different concentrations of ETP (50 μM, 100 μM, and 200 μM) to soybean plants. As we expected, plant height was decreased under both stress conditions (only WL or WL with ETP treatments) as compared to that of the control (Fig. 3A). When comparing among stress conditions, decreased plant height was observed in WL only and WL with 200 μM ETP application at 10 DAT and 15 DAT (Fig. 3A). Higher chlorophyll content was observed in the non-stress condition in all time periods (Fig. 3B). ETP-treated plants revealed gradually improved chlorophyll contents depending on the concentration of ETP as compared to that of plants with only WL stress (Fig. 3B).

**Fig. 3.**
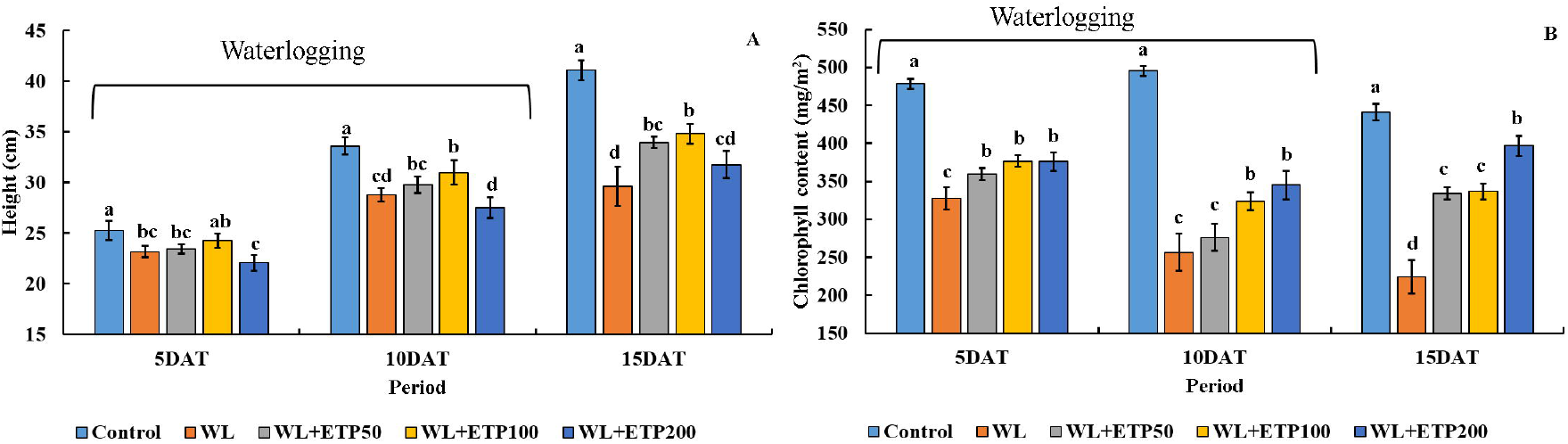
Influence of various concentrations of ethephon treatments on plant height and chlorophyll content. Soybean plants were exposed to WL for 10 days. Data were collected at 5 days intervals with three replications and the average ± standard error are shown (*n = 10*). Different letters indicate significant differences at *P < 0.05* and data were analyzed by Duncan’s multiple range test (DMRT). The abbreviation WL, DAT, and ETP mean WL, days after treatment, and ethephon, respectively.

#### Effects of ETP application on chlorophyll contents and fluorescence

To evaluate photosynthetic efficiency, we measured chlorophyll fluorescence at 15 days after WL treatment. According to our results, overall decreased OJIP values were observed in WL only and WL with ETP-treated soybean plants as compared to that of control plants (Fig. 4A). However, ETP-treated soybean plants showed a slightly improved OIJP value in comparison with that of WL-only treated soybean plants. In particular, sections from I to P exhibited large differences between ETP-treated soybean plants and WL-only treated soybean plants (Fig. 4A). Photosynthetic efficiency (Fv/Fm) also showed similar results to that of OJIP. Non-stressed soybean plants showed higher level of Fv/Fm (0.76) than other treatments. Comparison between WL and WL + ETP revealed improved photosynthetic efficiency in ETP-treated soybean plants (Fig. 4B).

**Fig. 4.**
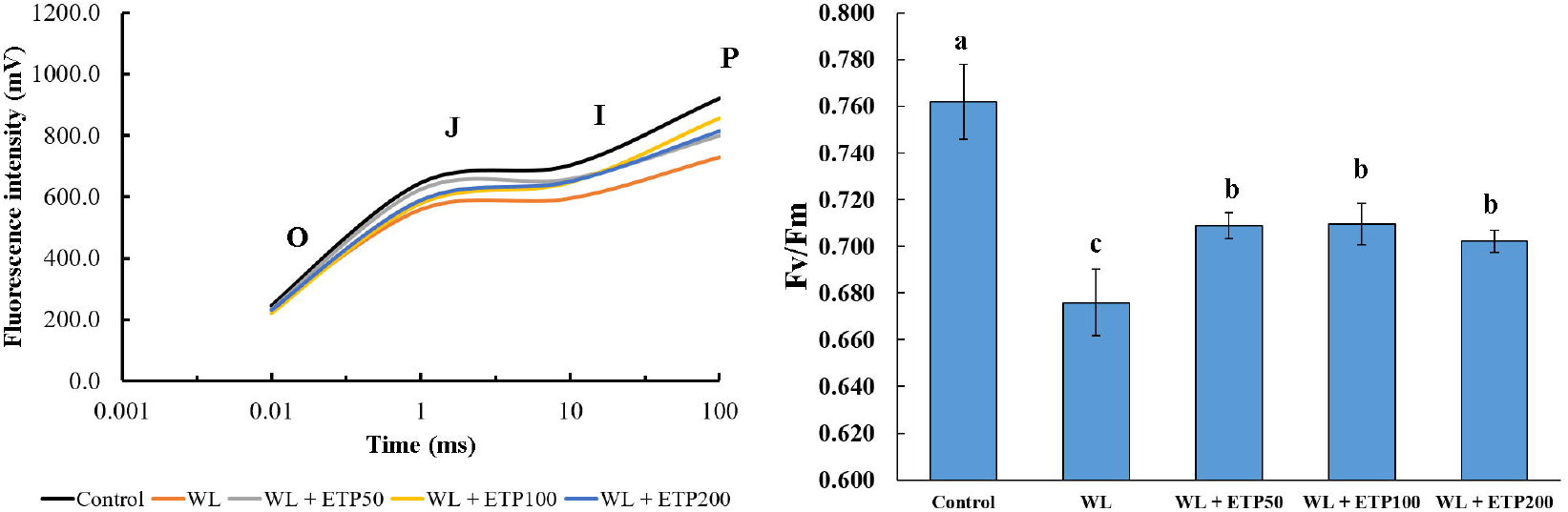
The OJIP parameters and photosynthetic efficiency (Fv/Fm) in soybean plants at 15 days after WL. Data were collected for three replications and the average ± standard error are shown (*n = 10*). The abbreviations WL, DAT, and ETP mean WL, days after treatment, and ethephon, respectively.

#### Influence of ETP application on endogenous plant hormones

We analyzed endogenous GA levels to elucidate physiological responses during stress periods. In higher plants, endogenous bioactive GAs (GA_1_, GA_3_, GA_4_, and GA_7_) are synthesized by two different pathways, one is the early 13-hydroxylation pathway and the other is the non-13-hydroxylation pathway (Kim *et al.,* 2016). According to a previous study, soybeans mainly produce bioactive GA_4_ through the non-13-hydroxylation pathway. Thus, we focused on determining the bioactive GA_4_ level and that of its intermediate precursor (GA_9_) and catabolite (GA_34_) (Kim *et al.,* 2015). In Fig. 5, GAs indicated are the sum of GA_4_, GA_9_, and GA_34_. GA contents exhibited relatively low levels in the control and ETP200 application as compared to other treatments, whereas GA contents were significantly increased in ETP50 and ETP100 in comparison with the WL-only treatment at 5 days after stress exposure. Enhanced GA levels were observed in ETP50 and ETP100 at 10 days of WL treatment (Fig. 5).

**Fig. 5.**
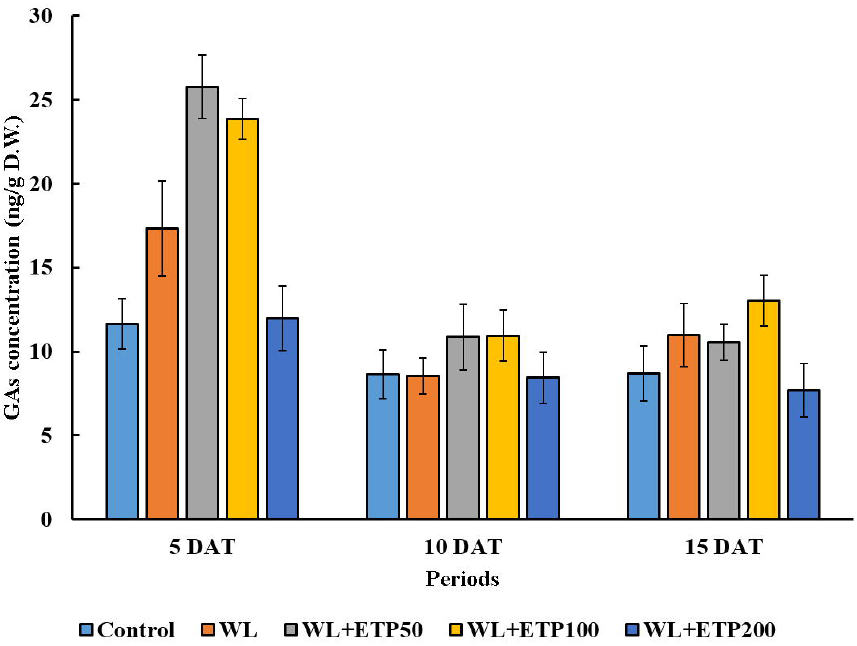
Influence of various concentrations of ethephon treatments on endogenous hormone levels. Soybean plants were exposed to WL for 15 days. Data were collected at 5-day intervals with three replications. In each figure, different letters indicate significant differences at *P < 0.05* and data were analyzed by Duncan’s multiple range test (DMRT) in the figure. The abbreviations WL, DAT, and ETP mean WL, days after treatment, and ethephon, respectively. In figure A, GA concentration means the sum of GA_4_, GA_9_, and GA_34_.

#### Influence of ETP application on macro element contents

The macro element contents are shown in Fig. 6. Carbon (C) content was higher in soybean plants as compared to other elements. Oxygen (O) content was increased when soybean plants were exposed to the WL condition. Similarly, enhanced P and Fe also occurred; the contents of both elements were increased in the WL condition in comparison with that of the control. However, overall element content was not higher (Fig. 6). However, N, K, and Ca contents showed different patterns. When we supplied the WL condition to soybean plants, N, K, and Ca contents were significantly decreased in WL and WL with ETP application as compared to that of the control (except for Ca content only in the WL-only treatment at 5 DAT). In particular, N and K content were significantly increased in ETP-treated soybean plants in comparison with WL-only soybean plants (Fig. 6). We could not detect any differences in H, Mg, or S content, although decreased total macro elements (sum of C, O, H, N, K, Ca, P, Mg, S, and Fe) content was observed in WL-only and ETP-treated soybean plants as compared to that of control at 10 DAT and 15 DAT (Fig. 6).

**Fig. 6.**
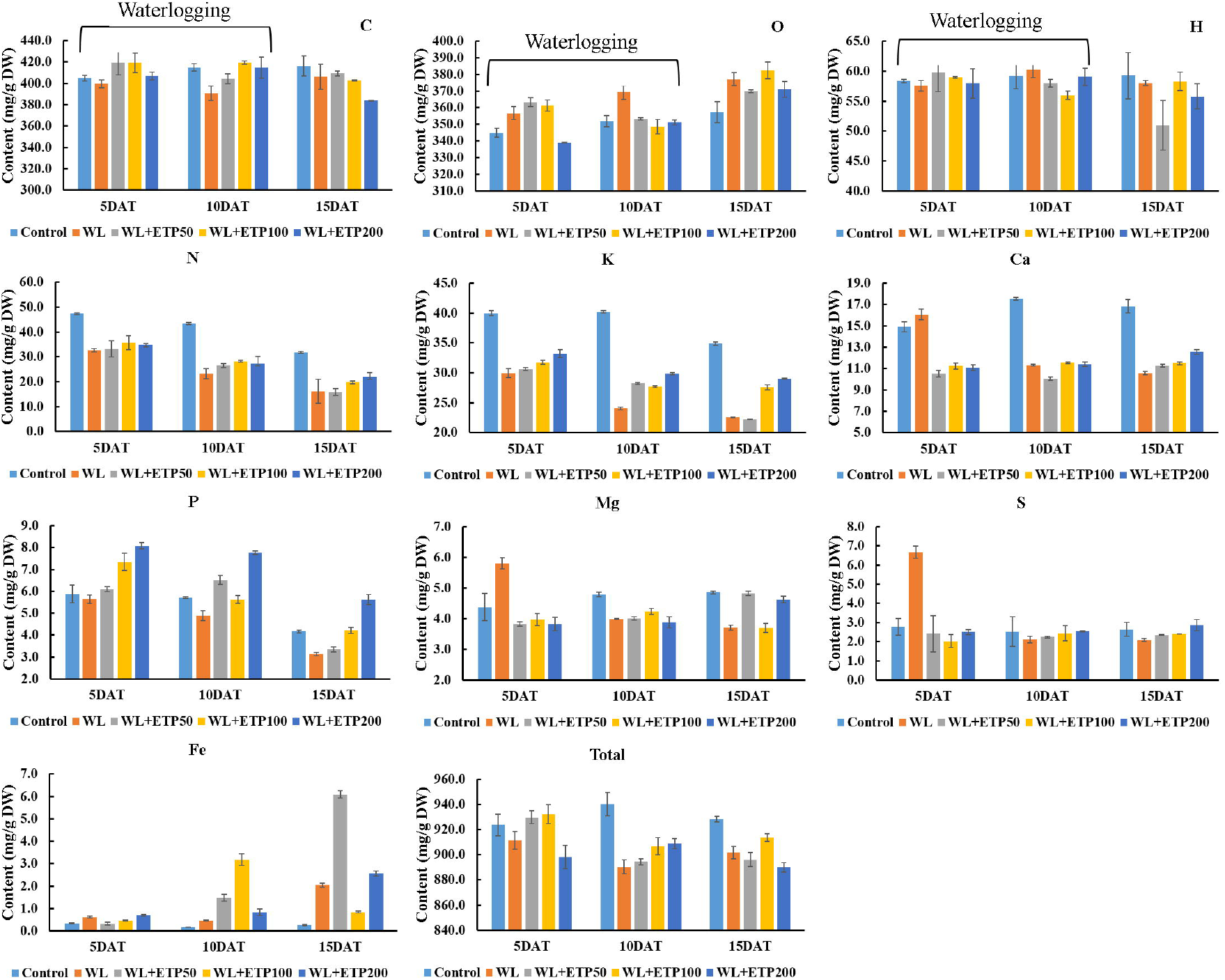
Mineral contents in soybean plants during and after WL condition. Data were collected at 5 days interval for 15 days and standard error was collected from two replications. The abbreviations WL, DAT and ETP mean WL, days after treatment, and ethephon, respectively. In the figure, total indicated sum of macro elements = C, O, H, N, K, Ca, P, Mg, S, and Fe.

#### Changes in amino acid contents

Methionine, proline, cysteine, and glutamic acids are known as an abiotic stress response. Methionine content was significantly decreased in soybean plants grown in WL conditions. However, increased methionine content was detected in all ETP-treated soybean plants in comparison with soybean plants grown in WL-only conditions (Fig. 7A). The same pattern was observed at all time points (5 DAT, 10 DAT, and 15 DAT). Proline content and that of glutamic acid also revealed that when soybean plants were exposed to WL conditions, proline and glutamic acid content was significantly decreased as compared to that of the control and the same tendency was observed at all time point (Fig. 7B, D). In ETP-treated plants, contents of proline and glutamic acid showed statistically similar or slightly increased values in comparison with the WL-only condition (Fig. 7B, D). Cysteine content did not exhibit a difference among treatments at 5 DAT. However, it was significantly decreased in WL and ETP-treated soybean plants (Fig. 7C). In comparison between WL-only and ETP-treated plants, cysteine content revealed a significant reduction at 10 DAT and no difference among treatments at 15 DAT (Fig. 7C). The sum of 17 amino acids (total amino acid) contents showed consistence result (Fig. 7E). Total amino acid contents were significantly decreased when soybean plants were exposed to the WL condition in comparison with that of the control plant, whilst concentration of amino acids was significantly improved with ETP application as compared to WL-only at 10 and 15 DAT (Fig. 7E).

**Fig. 7.**
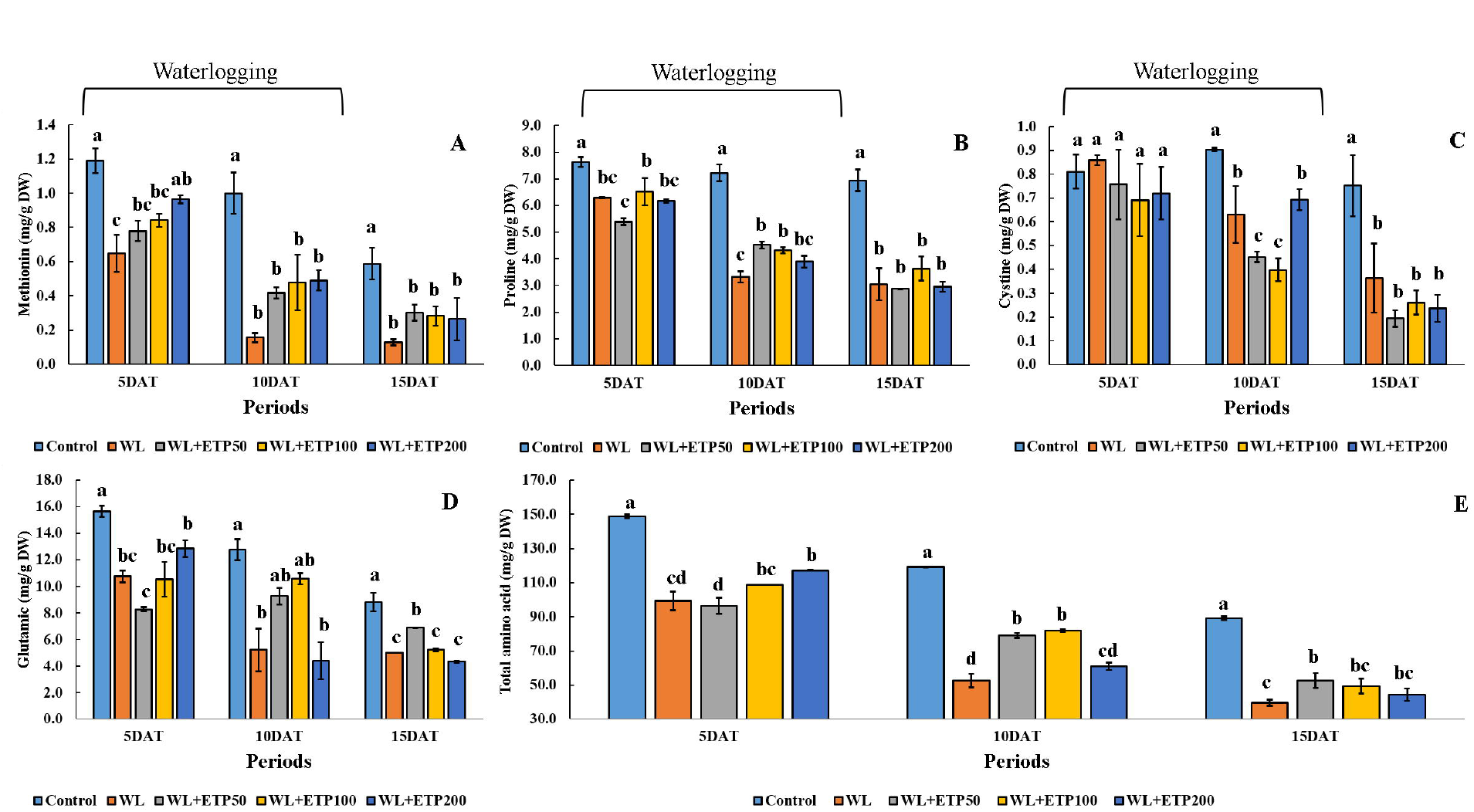
Amino acid contents in soybean plants during and after WL condition. Data were collected at 5 days interval for 15 days and standard error was collected from two replications. The abbreviations WL, DAT, and ETP mean WL, days after treatment, and ethephon, respectively. In the figure, total amino acid indicated sum of 16 amino acid = Ala, Arg, Asp, Cys, Glu, Gly, Ile, Leu, Lys, Met, Phe, Pro, Ser, Thr, Tyr, and Val.

#### Proteomics expression during WL conditions

To identify the protein expression pattern under WL conditions, we used soybean plants exposure to WL for 10 days. The 2-DE gel image showed a total of 63 different proteins in the expression pattern (Fig. 8). Among 63 proteins, we identified seven interesting proteins and the proteins were identified as ribulose-1, 5-bisphosphate carboxylase/oxygenase large subunit (RuBisCO) (Spot No. 615, 616, 1611, and 1702), trypsin inhibitor A (Spot No. 1002), glutathione s-transferase DHAR2 (Spot No. 1104), and glycoprotein (Spot No. 4101) (Table 1). Among the seven identified proteins, two proteins (Spot No. 615 and 616; red arrows) were up-regulated only in WL condition as compared to the control and WL with ETP treatments, whereas five proteins (Spot No. 1002, 1104, 1611, 1702, and 4101; orange and white arrows) were down-regulated in only the WL condition in comparison with the control and WL with ETP application (Fig. 8). Moreover, the expression of these proteins were recovered by 100 μM ETP application to soybean plants. Therefore, our data suggested that these proteins were participating in inducing resistance to WL stress.

**Fig. 8.**
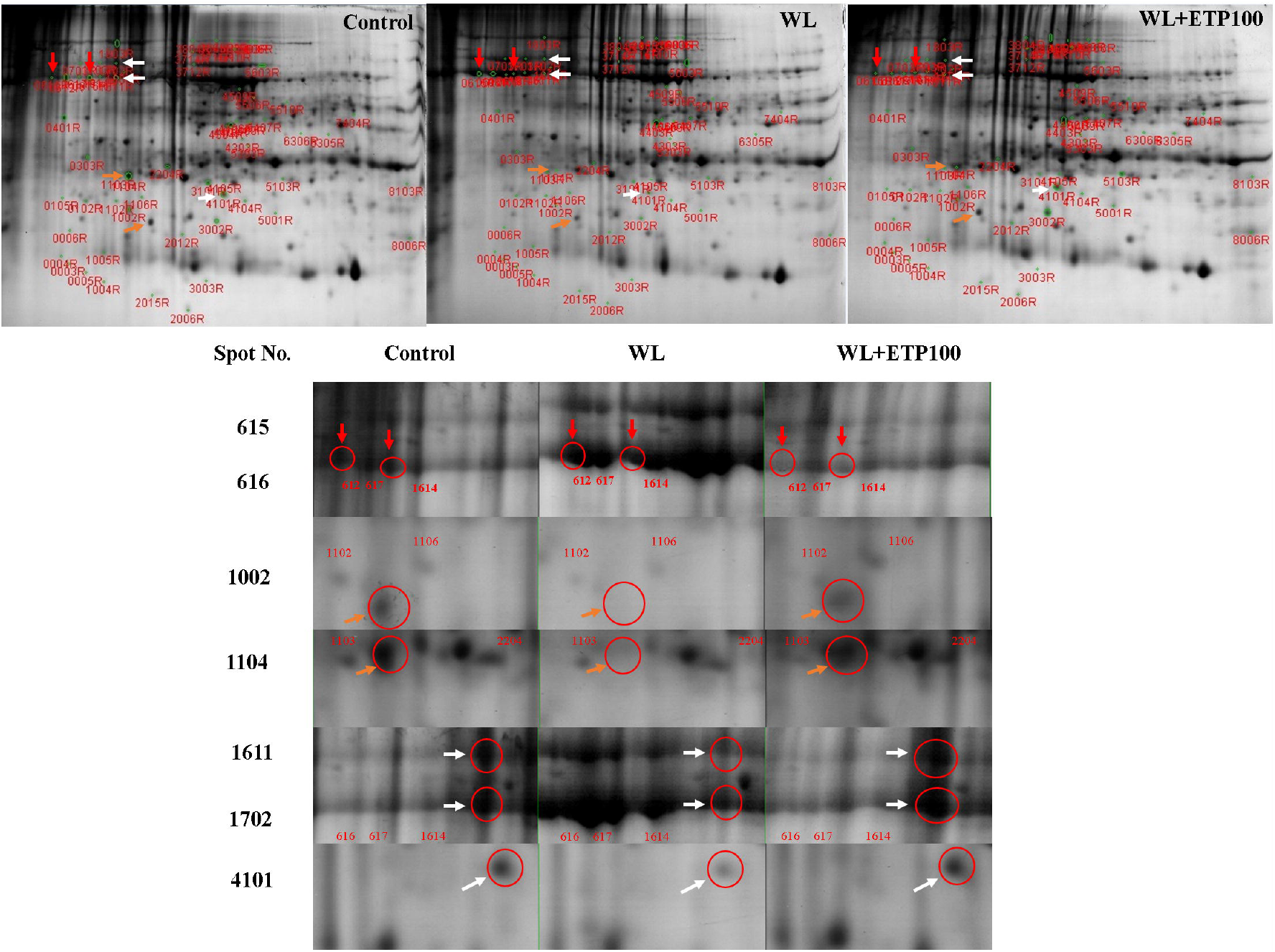
Root image and root surface area (RSA) in soybean plants during and after WL conditions. In the figure, white bar means 5 cm^2^ (W: 1 cm × L: 5 cm). Vertical bar with error bar is average with standard error (n = 5). In the same stress-exposure period, different letters indicate significant differences at *P < 0.05* and data were analyzed by Duncan’s multiple range test (DMRT) in the figure. The abbreviations WL, DAT, and ETP mean WL, days after treatment, and ethephon, respectively.

#### Root surface area (RSA)

To identify the ETP effect on soybean plants under WL condition, we analyzed RSA using the Flower Shape Analysis System software. As mentioned in the figure title, the white vertical bar indicates 5 cm^2^ [1 cm (W) × 5 cm (L)]. Adventitious roots did not appear in the control group of soybean plants, whereas they were very well developed in the WL with ETP treated soybean plants as compared to the WL-only treated soybean plants. The same results were observed in all time periods (Fig. 9). Root surface area (RSA) revealed that control soybean plants had significantly increased RSA as compared to WL and WL with ETP application at 5 DAT, 10 DAT and 15 DAT. RSA was increased in WL with ETP treated soybean plants as compared to WL-only treated soybean plants (Fig. 9). In particular, the application of ETP 100 and ETP 200 showed significantly increased RSA in comparison with ETP 50 at 15 DAT (Fig. 9).

**Fig. 9.**
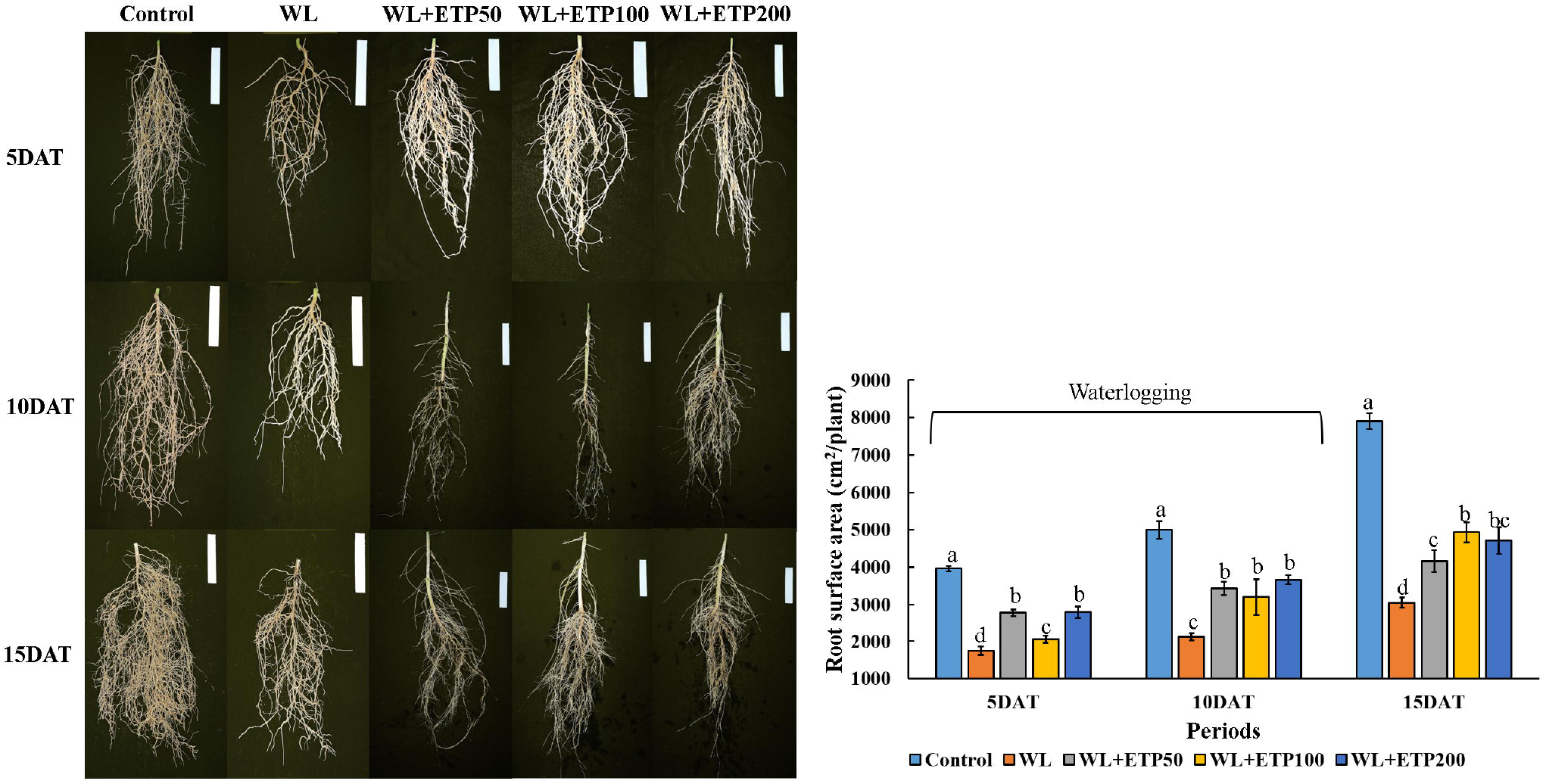
Protein expression levels in soybean plants. In the figure, orange and white color arrows indicated down-regulated proteins in the WL-only condition, whereas up-regulated proteins observed in WL in ethephon-treated plant. In addition, red color arrows point out highly up-regulated proteins under the WL-only condition.

#### Antioxidant activity and its mRNA expression level

According to our 2-DE results, glutathione s-transferase DHAR2 protein was down-regulated in the WL-only treated soybean plants, but it was recovered by ETP application. Thus, we measured activities of GSH and GR at the genetic and enzymatic level. GR activity of soybean shoots were decreased in the WL-only and WL with ETP-treated soybean plants as compared to that of the control plants (Fig. 10A). However, GR activity was increased in the WL with ETP-treated plants in comparison with the WL-only treated soybean plants (Fig. 10A). GR activity in the shoot revealed a similar pattern, whereas GR activity in the root did not show a regular pattern (Fig. 10C). Expression levels of *GmGR* exhibited differences between the shoot and root. In the shoot, the expression level of *GmGR* was lower in the WL-only and WL with ETP-treated soybean plants as compared to that of the control at 1 DAT (Fig. 11A). In comparison between WL-only and WL with ETP-treated plants, the expression level of *GmGR* was lower in WL with ETP-treated soybean plants than WL-only treated soybean plants, whereas the expression level of *GmGR* changed dramatically at 2 DAT. The greatest expression levels occurred for WL with 50 μM and WL with 100 μM ETP (Fig. 11A). In roots, the expression of *GmGR* was higher in WL-only treated plants than others, whereas WL with ETP-treated soybean plants exhibited a decreased expression level compared to that of the control and WL-only plants (Fig. 11D).

**Fig. 10.**
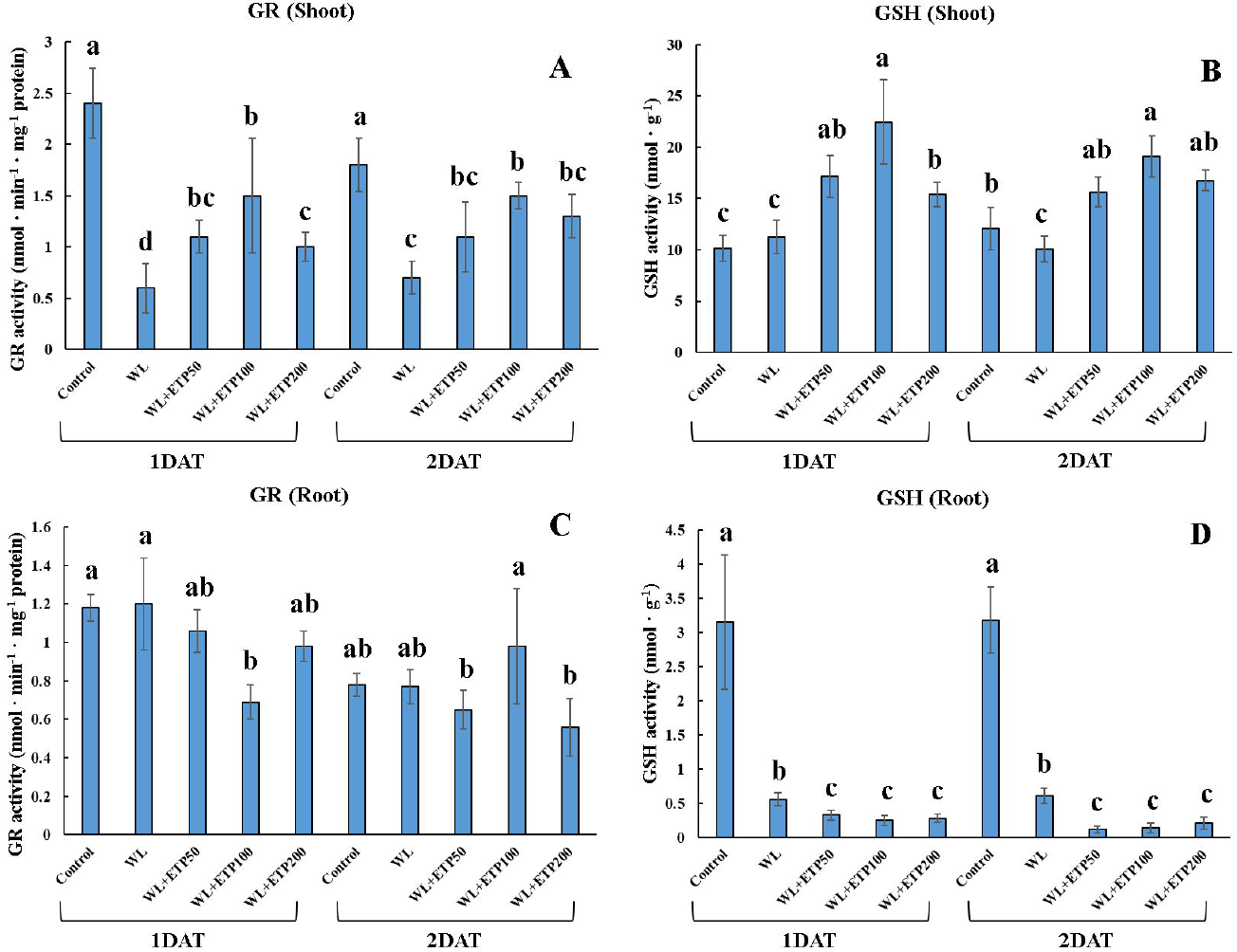
Influence of various concentrations of ethephon treatments on antioxidant (GR and GSH) activity. Soybean plants were exposed to WL for 2 days. Data were collected at 1 day intervals and in three replications. In each figure, different letters indicate significant difference at *P < 0.05* and data were analyzed by Duncan’s multiple range test (DMRT) in the figure. The abbreviations WL, DAT, and ETP mean WL, days after treatment, and ethephon, respectively.

The activity of GSH exhibited a significant difference between shoots and roots. In the shoot, GSH activity was significantly increased in WL with ETP-treated soybean plants as compared to that of the control and WL-only treated soybean plants; a similar pattern was observed in both stress exposure periods (Fig. 10B). However, GSH activity was significantly decreased in WL with ETP-treated soybean plants as compared to that of the control and WL-only treated soybean plants, and the same pattern repeated in both exposure periods (Fig. 10D). At the genetic level, GSH activity was very well described by *GmGST3* than *GmGST8* for both plant parts (shoots and roots) (Fig. 11B, C, E, F). WL with ETP-treated soybean plants exhibited a significantly increased expression level of *GmGST3* in the shoots, whereas the mRNA expression level of *GmGST3* was significantly lower in WL-ETP treated soybean roots as compared to that of other treatments (Fig. 11B, E). At 1 DAT, the expression level of *GmGST8* in the shoot revealed an increased tendency in WL-only and WL with ETP-treated plants compared with that of the control. However, we did not detect any significant difference between only WL and WL with ETP50 and WL with ETP100 (Fig. 11C). WLETP treated soybean plants exhibited a higher expression level of *GmGST8* as compared to that of WL-only and control plants in the shoots. In particular, WL with 50 μM ETP application showed the greatest expression level (Fig. 11C). In contrast, in the roots, expression levels of *GmGST8* were significantly decreased in WL with ETP-treated soybean plants as compared to that of the control (Fig. 11F). However, the expression level of *GmGST8* in the root did not exhibit any constant tendency for WL with ETP-treated soybean plants (Fig. 11F).

**Fig. 11.**
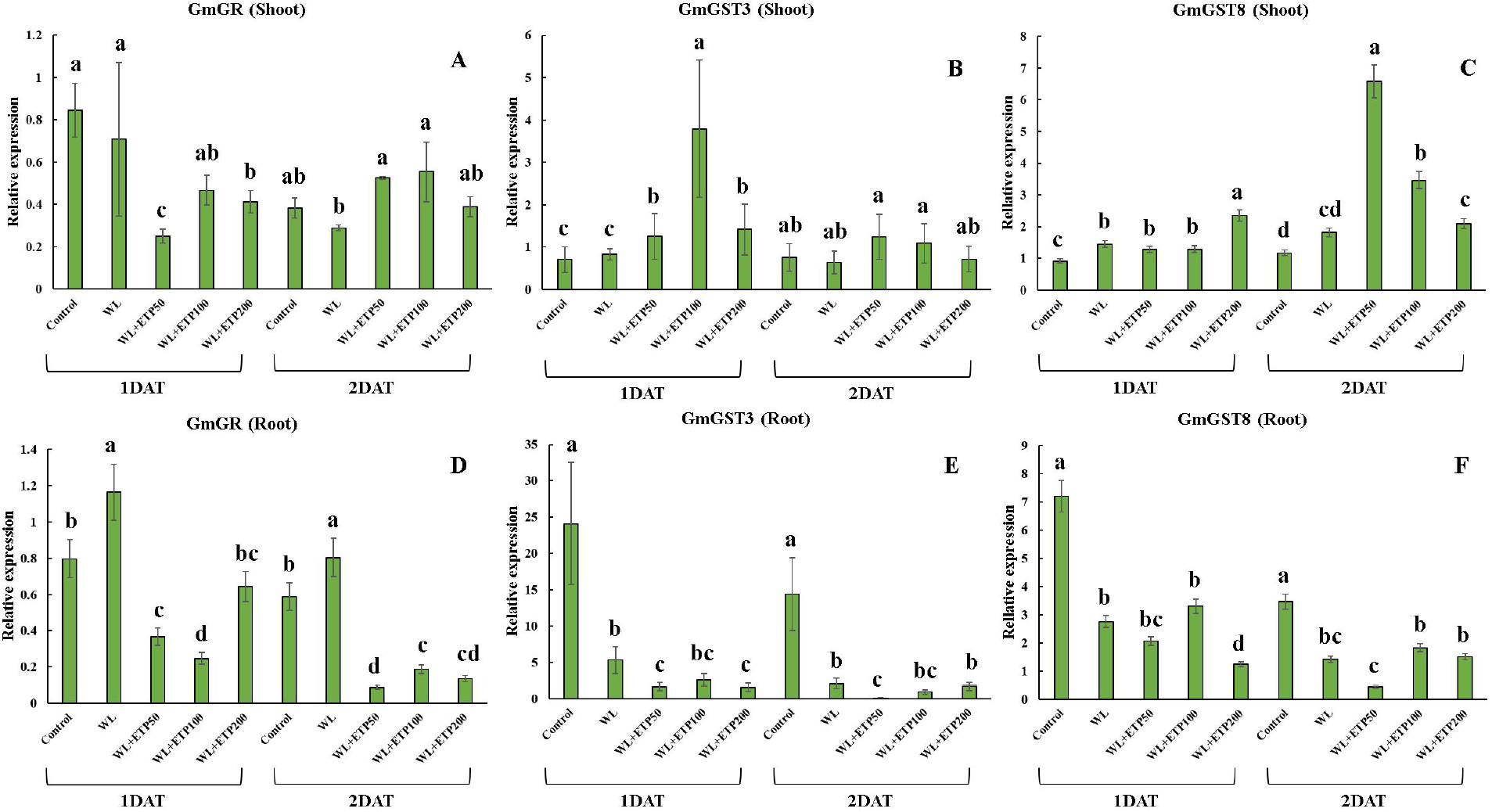
Influence of various concentrations of ethephon treatments on specific gene expression levels (GmGR, GmGST3, and GmGST8). Soybean plants were exposed to WL for 2 days. Data were collected at 1 day interval and was in three replications. In each figure, different letters indicate significant difference at *P < 0.05* and data were analyzed by Duncan’s multiple range test (DMRT) in the figure. The abbreviations WL, DAT, and ETP mean WL, days after treatment, and ethephon, respectively.

## Discussion

Plants are sessile and cannot escape from local changing environment. Thus, they are confronted with various abiotic and biotic stresses. Plants evolve different efficient defense related mechanisms to avoid stress conditions (Pérez-Clemente *et al*., 2012). According to Nishiuchi *et al*. (2012), plants under flooding conditions are faced with growth and developmental problems because of an inadequate supply of oxygen (Colmer and Flowers, 2008). Research related to flooding stress conducted in rice plants under submergence conditions (Hattori *et al*., 2009) showed that rice plants can respond via quiescence strategy or escape strategy. During quiescence strategy the rice plant saves carbohydrates during the submergence period. However, if plants defend stress conditions, preserved carbohydrates can be used for various physiological responses involved in growth and development (Schmitz *et al*., 2013). Whilst in escape strategy some rice cultivars exhibit rapid internode elongation to uptake oxygen with some parts above the water level (Hattori *et al*., 2009). Both strategies depend on ethylene-responsive transcription factors (Nishiuchi *et al*., 2012). Flooding tolerance mechanisms are very well known at the genetic level in rice plants. However, flooding mechanisms in soybean plants have not yet been fully elucidated. Previously, Nguyen *et al*. (2012) conducted QTL mapping by using contrasting soybean varieties against WL to identify flooding mechanisms in soybean plants. Several genes [*Sub1A*, *SNORKEL1 (SK1)* and *SNORKEL2 (SK2)*] related to flooding stress in rice plants were identified by QTL mapping. To date, several QTLs involved in WL resistance (Nguyen *et al*., 2012) have been reported. However, the QTLs were not identified by physiological mechanism or marker-assisted selection (Kim *et al*., 2015).

In the current study, significantly increased ethylene production was observed in a WL-tolerant variety; in addition, decreased ABA contents were detected. Based on previous research, we evaluated the effects of exogenous treatment of plant growth regulators in this study. According to our results (plant height, chlorophyll content, chlorophyll fluorescence), GA, KT, and ETP application caused improved resistance against WL in comparison with the control. Among three PGRs applied, we selected ETP as a candidate because GA application revealed WL mitigation via escape strategy (hyper-elongation of shoot). However, it would be difficult to use in the agricultural industry because all soybean plants showed lodging because of substantial internode growth (Seo *et al*., 2017). ETP-treated plants showed more enhanced shoot growth, as well as reduced leaf chlorosis in comparison with KT-treated soybean plants. Moreover, theoretically ETP application was closer to our previous findings (Kim *et al*., 2015). Thus, we selected ETP as the final candidate from among several PGR treatments.

To identify in detail the physiological and genetic differences between ETP-treated and control plants, we applied three different concentrations of ETP to soybean plants and measured several aspects of phenotypic variables, such as growth characteristics, endogenous hormones, and protein expression. Overall plant height showed an increased tendency in ETP-treated WL plants as compared to that of WL-only plants. Therefore, we needed to determine the tolerance mechanism in wetland plants to illustrate the difference between ETP-treated WL plants and WL-only plants. In rice plants, different resistance strategies were identified depending on the water level (Xu *et al*., 2006). Hattori *et al*. (2009) reported that increased shoot length was measured in a WL-tolerant rice variety under submergence conditions and the physiological response was induced by *SK1* and *SK2* genes. The *SK1* and *SK2* genes are included in the ethylene response factors (ERF). Thus, this gene accumulates GA or induces GA signal transduction to increase internode elongation. Finally, different shoot lengths were measured in deep water rice plants (Hattori *et al*., 2011). Similar results were reported in our previous research (Kim *et al*., 2015). According to Kim *et al*. (2015), significantly increased bioactive GA_4_ was observed in a WL-tolerant soybean variety. Therefore, based on the evidence above, increased shoot length in ETP application would be one of the stress avoidance strategies in rice plants and endogenous bioactive GA would participate in this response.

The photosynthesis in plants is primarily driven by two different photosynthetic apparatuses, including photosystem II (PSII) and photosystem (PSI) at the thylakoid membrane (Aro *et al*., 2005). The main role of PSII is oxidation of water to form oxygen and protons, followed by the transfer of protons to APT synthase for generating ATP (Bellafiore *et al*., 2005). The OJIP parameters indicate photosynthetic efficiency and have been broadly used for determining plant stress (Kalaji *et al.,* 2016). According to our results, when soybean plants were exposed to WL stress for 15 days, the OJIP parameter decreased in WL-only and WL with ETP-treated plants as compared to that of the control. Fv/Fm also showed a decrease in WL-only and WL with ETP-treated soybean plants. Therefore, we could have predicted that the WL condition induced a stress response, including photosynthetic efficiency in soybean plants. However, ETP-treated soybean plants during WL exhibited a smaller decrease for both the OJIP parameter and Fv/Fm value in comparison with that of WL-only treated soybean plants. Thus, we assumed that ETP supplementation to soybean plants during WL condition participated in stress mitigation.

In addition, among plant hormones, GAs are known as key signaling molecules involved in various physiological responses, such as seed germination, cell division, cell elongation, and stress responses (Kim *et al.,* 2016). In particular, GAs are deeply involved in the water stress escape strategy in rice. The up-regulation of various GAs is reported in rice plants during submergence and it helps the rice plant to expose some of its parts to the atmosphere because of hyper elongation in the rice stems. Finally, this physiological and morphological response confers resistance against submergence conditions in rice plants (Fukao *et al.,* 2011). Likewise, increased GA contents of WL-tolerant soybean cultivars were reported by Kim et al. (2015), who showed that endogenous GAs are involved in avoidance of WL stress because of the changes in GA levels to induce escape from excess water stress. In the current study, significantly increased GAs were detected in WL with ETP50- and ETP100-treated soybean plants. In particular, GA concentration was higher at relatively short-term (5 DAT) stress exposure. Therefore, our results suggested that exogenously applied ethylene participates in accumulation of endogenous GAs and this physiological phenomenon is one of the mechanism used to escape WL stress in soybean plants.

Most plants uptake primary inorganic ions such as nitrogen, calcium, potassium, and phosphorus through roots from the soil (Schachtman *et al.,* 1998). For that reason, root growth and development are very important to the absorbance of many elements. WL conditions in soybean plants is one of the restricted factors for root growth. When soybean plants were exposed to WL condition for 5 days, decreased root size was observed in the WL only condition but ETP treated soybean plants showed more increased root size as compared to only-WL condition. RSA was also decreased in the WL-only condition, whereas ETP-treated soybean plants showed increased RSA. Accordingly, WL conditions induced restricted root growth. However, exogenous application of ETP improved root growth. Consequently, we hypothesized that improved root growth because of ETP supplementation would result in increased element contents in soybean plants (Jackson *et al*., 2009). As we anticipated through the root images, total element contents were decreased in the WL-only condition. However, they were improved by ETP application. Thus, our results suggested that i) soybean root growth was inhibited under WL conditions. However, ETP supplementation induced improved root growth, ii) decreased the reduction in root condition under WL that causes decreased element uptake through soybean roots, which was thereby improved by ETP treatment, and iii) ETP application to soybean plants induced resistance against WL stress condition. Furthermore, we assumed that ETP application during WL conditions significantly involved K and P uptake among other elements because K and P contents were increased for all concentrations of ETP application in comparison with that of the WL-only condition.

Total amino acid contents were decreased by soil WL but they were increased in ETP-treated plants as compared to that of the WL-only condition. In particular, methionine content revealed significant differences between WL-only and WL with ETP-treated plants. ETP-treated soybean plants showed increased methionine content as compared to non-ETP treated soybean plants during WL conditions. WL tolerant soybean plants showed significantly increased endogenous ethylene production as compared to WL susceptible soybean plants, whereas significantly decreased methionine content was observed in WL tolerant soybean plants as compared to WL susceptible soybean plants. Thus, we assumed that WL tolerant soybean plants accumulated more endogenous ethylene to resist WL conditions. Therefore, less methionine content was observed in WL tolerant plants because methionine is a precursor of ethylene.

We induced high concentrations of endogenous ethylene production artificially in soybean plants to resist WL conditions. ETP application to soybean plants after WL, on the other hand, induced changes in the amino acid content in plants. We measured protein expression in different treatments and 63 different protein spots were observed. Among the 63 different spots, seven interesting protein spots were used for proteins identification. Among the seven proteins, two proteins (spot No. 615 and 616) were down-regulated in ETP-treated soybean plants as compared to control and WL-only treated soybean plants, and these were identified as RuBisCO proteins. However, the other two proteins (spot No. 1611 and 1702) were up-regulated in ETP-treated soybean plants as compared to WL-only treated soybean plants and they were also known as RuBisCO proteins. RuBisCO is located in chloroplasts in plants and participates in carbon fixation during photosynthesis (Schwender *et al.,* 2004). Therefore, this data indicated that ETP application to soybean plants during WL conditions participated in the photosynthetic process, in particular, in carbon fixation. However, the current data does not prove whether ETP application to soybean plants could up-regulate or down-regulate the activity of RuBisCO.

An addition three protein spots were identified as trypsin inhibitor A, glutathione stransferase DHAR2, and glycoprotein, and all three proteins were up-regulated in ETP-treated soybean plants as compared to WL-only soybean plants. Among them, glutathione stransferase DHAR2 is involved in reactive oxygen species (ROS) scavenging by participating in the ascorbate-glutathione cycle, as well as reactive nitrogen species (RNS) scavenging (producing GSNO) (Yun *et al.,* 2016). Therefore, we focused on the analysis of gene expression involved in activity of GSH (*GmGR* and *GmGSTs*) to elucidate genetic difference caused by ETP supplementation during WL conditions. The main role of GSH is a crucial process of detoxification and redox buffering in plants (Edwards *et al.,* 2000). GSH-dependent enzymes, glutathione S-transferases (GSTs), are included in plants for detoxification (Edwards *et al.,* 2000). Glutathione S-transferases (GSTs) are abundant proteins and are involved in xenobiotic detoxification, as well as acting as antioxidants by i) combining with oxidative degradation products, ii) acting as a glutathione peroxidase, and iii) removing lipid peroxides (Dalton *et al.,* 2009). Thus, we analyzed mRNA expression level of *GmGST3* and *GmGST8*. According to our results, expression levels of *GmGST3* were upregulated in all ETP-treated soybean plants (shoots) as compared to that of the control and WL-only plants. In addition, increased GSH activity was observed in ETP-treated soybean plants (shoots). In other words, this result indicated that ETP application to soybean plants during WL could stimulate up-regulation of GST3 expression. Therefore, increased GSH activity, up-regulated proteins, and enhanced glutathione s-transferase DHAR2 were observed. Therefore, we assumed that all of these sequences of physiological responses were induced to mitigate WL conditions in soybean plants because of the exogenously applied ethylene producer.

## Acknowledgments

This work was carried out with the support of the Cooperative Research Program for Agriculture Science & Technology Development (Project No. PJ010990) Rural Development Administration, Republic of Korea.

## Supplementary data

**Supplementary Table 1.** Information on plant growth regulators (PGRs) and application concentration.

**Supplementary Table 2.** GC-MS-SIM and HPLC conditions for endogenous GA analysis.

**Supplementary Table 3.** Primer sequences for qRT-PCR

